# Heart enhancers with deeply conserved regulatory activity are established early in development

**DOI:** 10.1101/312611

**Authors:** Xuefei Yuan, Mengyi Song, Patrick Devine, Benoit G. Bruneau, Ian C. Scott, Michael D. Wilson

## Abstract

During the phylotypic period embryos from different genera show similar gene expression patterns, implying common regulatory mechanisms. To identify enhancers involved in the initial events of cardiogenesis, which occurs during the phylotypic period, we isolated early cardiac progenitor cells from zebrafish embryos and characterized 3838 open chromatin regions specific to this cell population. Of these regions, 162 overlapped with conserved non-coding elements (CNEs) that also mapped to open chromatin regions in human. Most of the zebrafish conserved open chromatin elements tested drove gene expression in the developing heart. Despite modest sequence identity, human orthologous open chromatin regions could recapitulate the spatial temporal expression patterns of the zebrafish sequence, potentially providing a basis for phylotypic gene expression patterns. Genome-wide, we discovered 5598 zebrafish-human conserved open chromatin regions, suggesting that a diverse repertoire of ancient enhancers is established prior to organogenesis and the phylotypic period.

## Introduction

The developmental hourglass model predicts a phylotypic stage during mid-embryogenesis when species within the same phylum display the greatest level of morphological similarities^1,2^. The hourglass model is also supported by comparative transcriptomic studies that demonstrated the most conserved gene expression occurred at the phylotypic stage^3–5^. The idea that conserved phylotypic gene expression is established through conserved enhancers is supported by several comparative epigenomic studies^6–10^. While most molecular studies of the phylotypic period have focused on whole embryos, recent evidence suggests that the exact developmental timing of maximal conservation varies in a tissue-specific manner^8^. We are only beginning to understand how conserved transcriptional programs for individual developmental lineages are set up prior to the phylotypic stage.

The heart, derived from the mesoderm, is the first organ formed during embryogenesis. Heart development is orchestrated by conserved cardiac transcription factors (TFs) binding to cis-regulatory elements (CREs)^11,12^. Crucial cardiac specification events occur during early embryogenesis^13–16^. For example, distinct subtypes of mouse cardiac progenitors emerge within the gastrula stage preceding the expression of the canonical cardiac progenitor marker *Nkx2*.*5*, long before any organ structure is formed^13–15^. However how this potential early cardiac specification is controlled by enhancer elements, and the extent to which this process is evolutionarily conserved, is not known.

Heart enhancers have been extensively characterized in studies utilizing genome-wide profiling techniques including chromatin immunoprecipitation followed by DNA sequencing (ChIP-seq)^17–22^, computational predictions^23,24^ and *in vivo* validation in mouse and zebrafish embryos^17,18,20,23,24^. To date, the majority of *in vivo* heart enhancer discovery and validation experiments were performed in embryos, when the heart chamber structures were already established, or in adult hearts^17,18^. *In vitro* differentiation of embryonic stem cells (ESCs) into cardiac progenitors have also yielded insights into early cardiac development and have enabled the discovery of cardiac enhancers^19,21,22^. However, more work is needed to identify the CREs that regulate the early differentiation of mesoderm progenitors to cardiac lineages in the context of the developing embryo.

Despite the highly conserved cardiac TFs necessary for heart development, heart enhancers identified at mouse E11.5 show limited phylogenetic conservation compared to brain enhancers identified at the same developmental stage^8,17^. However, analysis of putative enhancers in mesoderm cells, derived from embryonic stem cells, show higher evolutionary constraint than the enhancers identified after organogenesis^8^. This suggests that the regulatory elements that establish the conserved cardiac transcriptional program may exist at the initial stages of heart development, presumptively during the time window from naïve mesoderm to cardiac progenitors.

Here, we set out to discover enhancers that are active in cardiac progenitor cells prior to the expression of the cardiac progenitor marker *Nkx2*.*5*. We generated a zebrafish GFP reporter line driven by a mouse *Smarcd3* enhancer (*Smarcd3*-F6) that is active in early gastrulating mesoderm^13^, in order to enrich for early zebrafish cardiac progenitors. We profiled gene expression and the accessible chromatin landscape of GFP positive cells using the assay for transposase-accessible chromatin using sequencing (ATAC-seq), single cell mRNA-seq, lineage tracing, and RNA *in situ* hybridization, both in wildtype embryos and following knockdown of the essential cardiac transcription factors Gata5/6. Results from these experiments indicated that we had purified a population of cells enriched for cardiac progenitors. Using direct and indirect DNA alignments^25,26^, we identified accessible chromatin regions shared between zebrafish and human. We found that these conserved accessible chromatin elements were highly associated with developmental transcription factors that are regulated by Polycomb repressive complex 2 (PRC2). We confirmed the cardiac activity and functional conservation of many anciently conserved open chromatin regions using *in vivo* reporter assays. In sum, our study identified a set of conserved cardiac enhancers established prior to the phylotypic period, before the heart and other organ primordia appear, potentially providing a basis for common gene expression patterns across genera. Furthermore we uncovered ~6000 anciently conserved open chromatin regions that likely serve as enhancers for other cell lineages.

## Results

### The mouse *Smarcd3*-F6 enhancer marks cardiac progenitors in zebrafish

To examine the earliest events that contribute to cardiac lineage specification in zebrafish, we needed a means to isolate early cardiac progenitor cells *in vivo*. To identify a zebrafish marker that could drive GFP expression in cardiac progenitor cells prior to *nkx2*.*5* expression, we tested a recently described early mouse cardiac enhancer, *Smarcd3*-F6^13^ in our zebrafish model. Lineage tracing experiments demonstrated that this enhancer labeled cardiac progenitor cells prior to *Nkx2*.*5* expression in mouse embryos^13^. We also found that the *Smarcd3*-F6 region was enriched for the active enhancer mark H3K27ac and contains a CRE co-bound by several conserved cardiac TFs (GATA4, NKX2.5, TBX5) in mouse ESC differentiated cardiac precursors (CP) and cardiomyocytes (CM)^19,22^ (Supplementary Fig. 1A).

To test if the *Smarcd3*-F6 enhancer functions as an early marker of cardiac progenitors in zebrafish, we generated a *Tg(Smarcd3-F6:EGFP*) transgenic line (Fig. 1A). RNA *in-situ* hybridization against *gfp* showed the enhancer activity was robustly detected at 6 hours post-fertilization (hpf) along the embryonic margin (Fig. 1B), which contains mesendodermal progenitors including future cardiac cells^27^. Over the course of gastrulation, GFP positive cells migrated to encompass positions in the <u>a</u>nterior and <u>p</u>osterior lateral <u>p</u>late <u>m</u>esoderm (ALPM and PLPM) (Fig. 1B). Co-immunostaining comparing *Tg(Smarcd3-F6:EGFP*) and *Tg(nkx2.5:ZsYellow*) expression indicated that the *Smarcd3*-F6 enhancer marked almost all cardiac mesoderm expressing *nkx2.5* at early somite stages (13 hpf) (Fig. 1C). *Tg(Smarcd3-F6:CreERT2*) lines were generated to trace the fate of *Smarcd3*-F6 labelled cells. By crossing *Tg(Smarcd3-F6:CreERT2*) to a *Tg(βactin2-loxP-dsRed-loxP-GFP*) reporter line, we found that following 4-hydroxytamoxifen (4-HT) addition at 8 hpf, cells labeled by the *Smarcd3*-F6 enhancer contributed to heart formation in later development (Supplementary Fig. 1). Although the *Smarcd3*-F6 enhancer shows limited mammalian, and no zebrafish sequence conservation (Supplementary Fig. 1A), the early labeling of cardiac lineages in zebrafish indicated that this enhancer would allow us to isolate a cell population enriched for cardiac progenitors.

**Figure 1.**
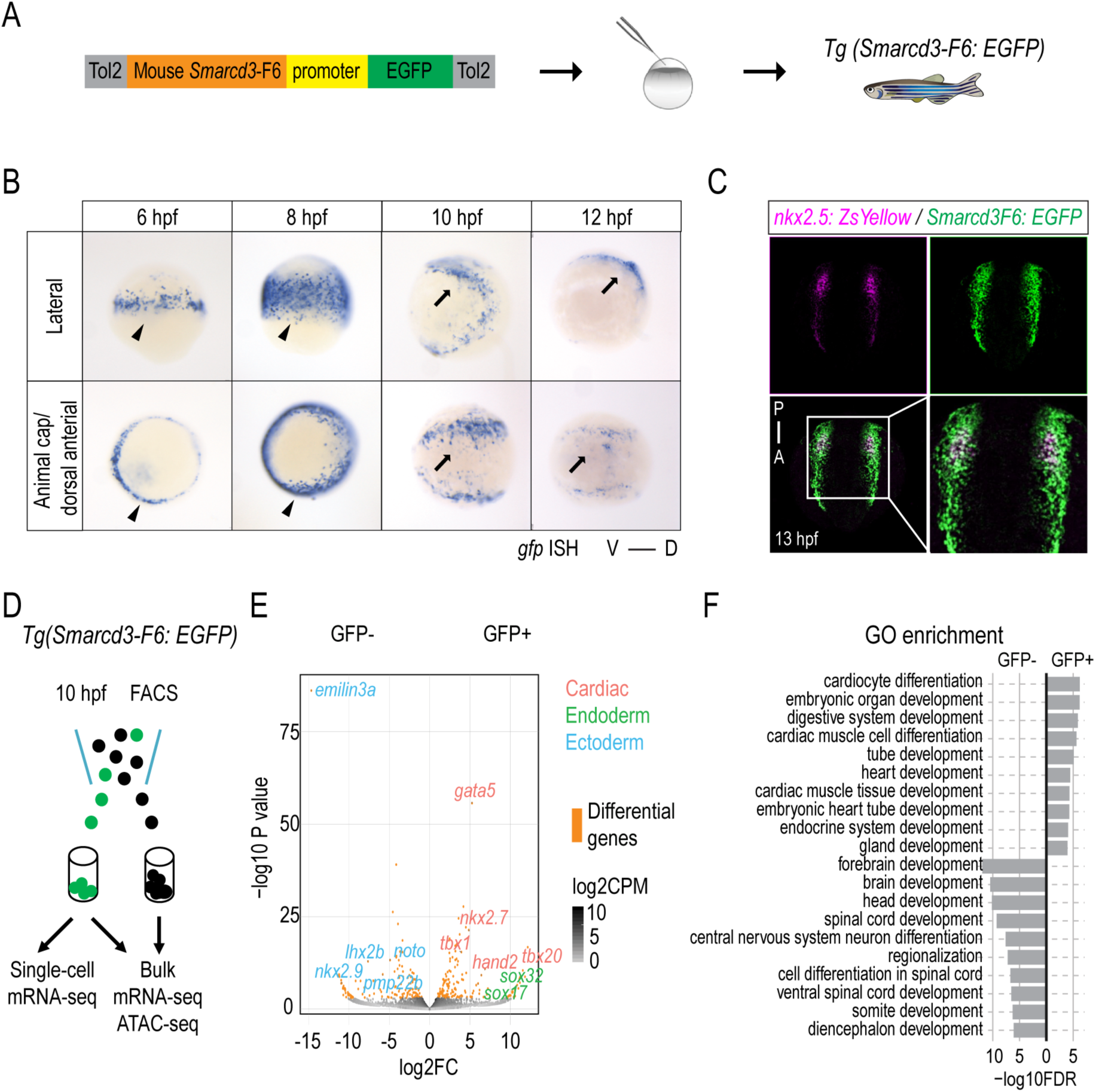
Mouse *Smarcd3*-F6 enhancer labels early cardiac progenitors in zebrafish. (A) Generation of *Tg(Smarcd3-F6:EGFP*) zebrafish line (B) *In-situ* hybridization against *gfp* transcripts on *Tg(Smarcd3-F6:EGFP*) transgenic embryos. *Smarcd3-F6* enhancer marks lateral margins (arrow heads) during gastrulation and ALPM regions (arrows) after gastrulation. (C) Immunostaining of ZsYellow and GFP on *Tg(Smarcd3-F6:EGFP*) and *Tg(nkx2.5:ZsYellow*) double transgenic embryos. Cells expressing ZsYellow were marked by GFP as well. (D) Workflow of mRNA-seq and ATAC-seq experiments. (E) Volcano plot showing genes differentially expressed between *Smarcd3-F6*:GFP+ and *Smarcd3-F6*:GFP- populations (FDR < 0.05, log2FC >1). (F) Top 10 most enriched GO terms obtained from genes that were significantly more highly expressed in *Smarcd3-F6*:GFP+ and *Smarcd3-F6*:GFP- populations.

To further characterize the population marked by the *Smarcd3*-F6 enhancer, we conducted bulk mRNA-seq and ATAC-seq at 10 hpf on *Smarcd3*-F6 labeled (GFP+) and unlabeled (GFP-) cells (Fig. 1D). mRNA-seq revealed 316 genes differentially expressed between the GFP*+* and GFP- population (FDR < 0.05, log2FC >1). Known cardiac (*gata5*, *nkx2.7*, *tbx20*, *hand2*) and endoderm (*sox32*, *sox17*) markers showed significantly higher expression in *Smarcd3*-F6 labelled cells while ectoderm genes were relatively depleted (Fig. 1E). Consistent with these results, genes showing higher expression in the GFP+ population displayed enrichment for processes related to cardiovascular and endoderm development, whereas genes enriched in GFP- cells were enriched for those involved in nervous system development (Fig. 1F).

To further dissect the putative cardiac progenitors within the 10 hpf *Smarcd3*-F6 labeled population, we performed single-cell mRNA-seq on 96 GFP+ cells. The average of the assembly of all single-cell transcriptomes correlated well with the bulk mRNA-seq results (R=0.93) (Supplementary Fig. 2A). Unsupervised clustering using genes differentially expressed between GFP+ and GFP- populations grouped cells into three broad clusters, which represented putative endodermal (*gata5^+^*, *sox17^+^*, *sox32^+^*), mesodermal (*gata5^+^*, *sox17^-^*, *sox32^-^*) and ectodermal (*gata5^-^*, *sox2^+^*, *sox3^+^*) populations (Supplementary Fig. 2B). Within the mesoderm cluster, we could identify a potential cardiac subgroup co-expressing known cardiac genes *gata5*, *hand2*, *tbx20*, and *tmem88a* (Supplementary Fig. 2B,C). Together, our transcriptome analyses demonstrated that cells labeled by the *Smarcd3*-F6 enhancer are enriched for cardiac lineages, with a putative cardiac progenitor population apparent by 10 hpf.

### Open chromatin regions enriched in the *Smarcd3*-F6 labeled population

Open chromatin profiles often identify the genomic regions where TFs and their co-factors bind and function^28,29^. Using ATAC-seq, we detected 155,879 open chromatin regions (ATAC-seq peaks) in GFP+ population and 153,777 in GFP- population. Our ATAC-seq peaks overlap with markers of active promoters (H3K4me3) and enhancers (H3K27ac, H3K4me1) that were previously identified from ChIP-seq experiments on whole embryos of similar stages^6^, as well as additional genomic regions, likely due to purifying a small subset of cells away from the bulk embryo (Supplementary Fig. 3A). Most ATAC-seq peak regions (n=195,466) shared similar ATAC-seq signals in both GFP+ and GFP- populations. 5,471 peaks showed significant quantitative differences (Fig. 2A) with 3,838 peaks specifically increased in *Smarcd3*-F6 labeled cells (‘GFP+ specific’), and 1,633 peaks specifically increased in unlabeled cells (‘GFP- specific’).

**Figure 2.**
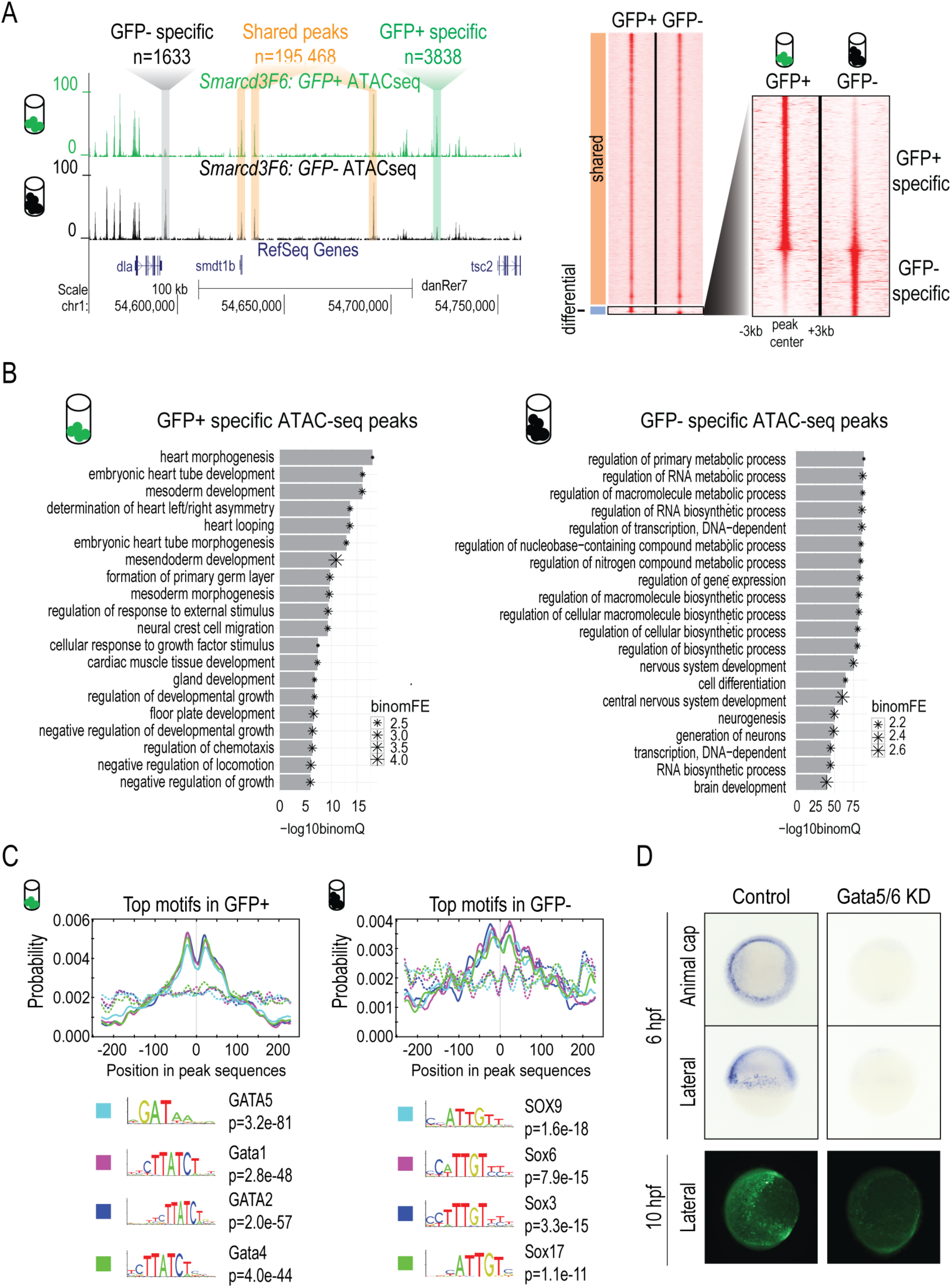
Open chromatin landscape of *Smarcd3*-F6 labeled population. (A) Genome browser view and heatmap showing ATAC-seq peaks shared between GFP+ and GFP- populations or enriched in one population. In the heatmap, read intensity for regions within 3kb of the peak center was plotted for each peak. (B) Barplot showing the 20 most enriched GO biological function terms obtained from GFP+/- specific peaks using GREAT analysis. (C) Probabilities of the top 4 enriched motifs within the GFP+/- specific peaks calculated by CentriMo. Each curve shows the probability of the best match to a given motif occuring at a given position in the input sequences. Solid lines represent probabilities caculated from query peak sets (GFP+/- specific peaks) while dash lines show that from the background sequences (shared peaks). (D) GFP *in-situ* and immunostaining on *Tg(Smarcd3-F6: EGFP*) emrbyos that were uninjected (control) or injected with *gata5/6* morpholinos. All staining and imaging were performed under the same condition for the control and *gata5/6* KD groups.

Overall both the GFP+ specific and GFP- specific open chromatin regions were depleted for proximal promoter regions and enriched for introns and intergenic regions (Supplementary Fig. 3B). Using the enrichment tool GREAT^30^, we found that the GFP+ specific open chromatin regions were enriched for heart development-related processes, based on proximity to genes (heart morphogenesis: FDR 1.09e-18, embryonic heart tube development: FDR 8.91e-17, binomial test, Fig. 2B). GFP- specific open chromatin regions were enriched for metabolic, gene expression and neural development related processes (regulation of primary metabolic process: FDR 5.60E-89, regulation of transcription, DNA-dependent: FDR 1.40E-86, nervous system development: FDR 1.35E-75, binomial test, Fig. 2B). Overall, the transcriptional profiles and open chromatin regions enriched in the GFP+ and GFP- specific populations further indicate that the *Smarcd3*-F6 enhancer marks cardiac progenitor cells.

We then asked which TF motifs were overrepresented in GFP+ specific peaks. We found GATA motifs showed the strongest enrichment, consistent with the crucial roles GATA factors play in heart and endoderm development^31–34^ (Fig. 2C). In contrast, GFP- specific peaks were most highly enriched for SOX motifs (Fig. 2C). Like *Gata4* and *Gata6* in mice^35–40^, *gata5* and *gata6* play redundant but critical roles for zebrafish heart formation^31,33^. To test if the activity of the *Smarcd3*-F6 enhancer is regulated by Gata5 and Gata6 in zebrafish, we performed *gata5* and *gata6* knock-downs by injecting morpholinos into *Tg(Smarcd3-F6: EGFP*) embryos (Fig. 2D). Supporting our motif enrichments we found that GFP expression by *Smarcd3*-F6 enhancer requires Gata5 and Gata6.

### Comparative epigenomic analysis of accessible chromatin elements reveals phylogentically conserved cardiac enhancers

To identify regions of open chromatin that are conserved between zebrafish and human, we used two well-defined conserved non-coding element (CNE) datasets, zCNE^25^ and garCNE^26^. Both of these two resources contain conserved regions identified using direct alignment and indirect homology bridged by intermediate species (Fig. 3A). We associated zebrafish ATAC-seq peaks to CNEs if they overlapped a zebrafish-human or zebrafish-mouse CNE. Most accessible chromatin-associated CNEs (~70%) were fully contained within ATAC-seq peaks. On average 30% of the length of these ATAC-seq peaks were comprised of CNEs. Within a total of 200,937 ATAC-seq peaks, we found 6294 (3.1%) or 6047 (3.0%) shared sequence conservation with human or mouse respectively (see Methods, Supplementary Fig. 3C and Supplementary Table 1 for details). Of these 6294 zebrafish-human ATAC-seq peaks 176 were GFP+ specific, 264 were GFP- specific.

**Figure 3.**
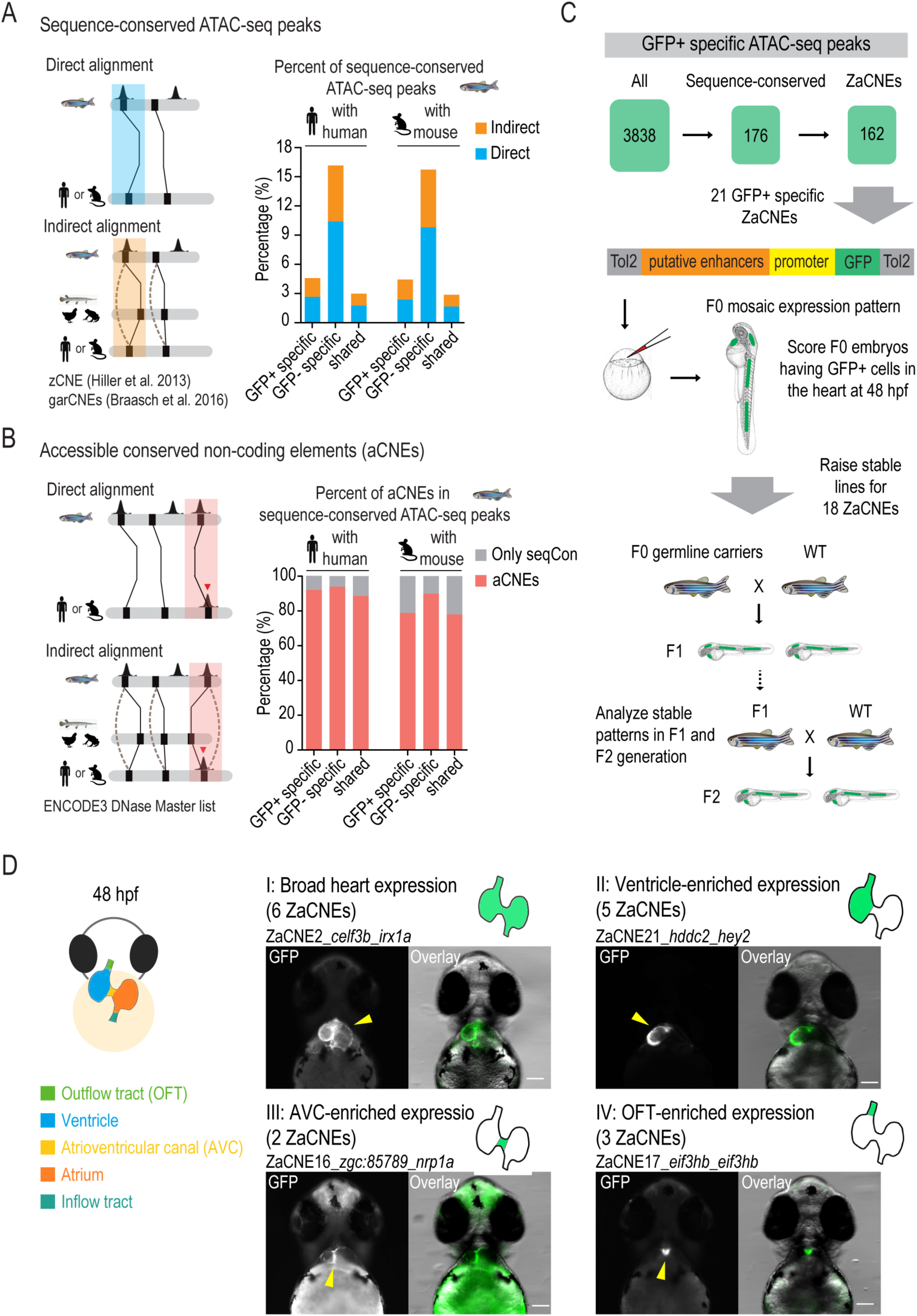
dentification and *in vivo* characterization of accesible conserved non-coding elements (aCNEs). (A) Bar plot showing the percentage of conserved regions in all GFP+/- specific and shared ATAC-seq categories. Solid lines in the cartoon indicate direct sequence alignment and dotted lines show cases of indirect alignment bridged by intermediate species or ancestry reconstruction. (B) Bar plot showing the aCNEs/sequenced conserved (seqCon) percentage. Most seqCon open chromatin regions were identified as having shared orthologous accessible chromatin. (C) Strategy for identifying and characterizing GFP+ specific aCNEs in zebrafish. Stable transgenic lines for 18 aCNEs were characterized. (D) Cartoon showing the zebrafish heart anatomy at 48 hpf (left panel). Heart expression patterns observed in ZaCNEs transgenic lines were classified into 4 categories (right panel), including I: broad expression in the whole heart; II: ventricle-enriched expression; III: atrioventricular (AVC)-enriched expression; IV: outflow tract (OFT)-enriched expression. One representative example of each category was shown and the numbers of ZaCNEs fall into each category were indicated in the brackets. The two closest genes near each ZaCNEs were indicated. All images were taken from the ventral view and all scale bars represent 100 μm.

We found that GFP- specific peaks were ~4 times more enriched for conserved regions than the GFP+ ones (*P*=1.40e-46 using human CNEs; Chi-square test) (Fig. 3A). Previous work has shown that 30-45% of forebrain, midbrain and limb enhancers overlapped regions of extremely high sequence constraint, in contrast to only ~6% of cardiac enhancers17. Given that GFP+ and GFP- populations were enriched for cardiac and brain lineages respectively (Fig. 1F), our observations were consistent with these previous findings^17,41^.

Comparing genomic features (e.g. TF binding, chromatin accessibility, histone modifications) between species is a potentially powerful way to identify conserved regulatory function within alignable sequence for specific tissues or cell types^42–47^. For ATAC-seq peaks overlapping CNEs, we asked if the orthologous human CNEs also contained accessible chromatin according to DNase I hypersensitivity sites (DHS) reported by ENCODE3^48^ (Fig. 3B). The majority of the zebrafish ATAC-seq peaks overlapping zebrafish-human or zebrafish-mouse CNEs overlapped human or mouse CNEs containing accessible chromatin (89% for human (5598/6294) and 79% (4747/6047) for mouse; Fig. 3B,C). Overall ~3% of the total zebrafish ATAC-seq peaks were identified as accessible non-coding regions that are shared between zebrafish and human, or zebrafish and mouse. We refer to conserved accessible chromatin connected through CNEs as aCNEs.

We asked whether any genomic features could distinguish aCNEs from bulk open chromatin regions. First, we found conserved ATAC-seq peaks had a wider boundary (average 706 bp compared to 475 bp, P value < 0.001; permutation test), stronger open chromatin signals and slightly higher GC content (average 46% compared to 44%, P value < 0.001; permutation test) than the bulk zebrafish ATAC-seq peaks (Supplementary Fig. 3D,E). On average, one zebrafish aCNE has 1.6 human and 1.4 mouse orthologous aCNEs based on the open chromatin data we used (Supplementary Fig. 3C, see methods for details.)

We compared our aCNEs to ultraconserved non-coding elements in the human genome^49^, which were defined as sequence elements with no mismatches for at least 200 base pairs between orthologous regions in the human, rat and mouse genomes. While 30% of ultraconserved elements overlap human aCNEs, only 3% of the aCNEs overlap ultraconserved elements (Supplementary Table 2). The fact that nearly 40% (2228/5598 for zebrafish-human aCNEs) of the aCNEs were found using indirect alignments indicates that many of these ancient aCNEs show limited sequence conservation and would likely have been overlooked using standard multiple genome alignment methods.

### Conserved accessible chromatin regions drive expression in the developing heart

To gain insight into the enhancer activity of aCNEs, we compared them to elements tested in the VISTA enhancer database, which is the most comprehensive database of functionally validated enhancers. Roughly 10% (556/5849) of human or mouse aCNEs have been experimentally tested *in vivo* and reported in the VISTA Enhancer Browser, two thirds of which were recorded as positive enhancers (2017/07/29 release; https://enhancer.lbl.gov/). 43/162 of the GFP+ aCNEs that we found overlapped heart enhancers that were predicted using curated epigenomic data^23^.

To determine if some conserved open chromatin elements are bound by cardiac TFs during heart development, we collected published ChIP-seq and ChIP-exo data for GATA4, NKX2.5, TBX5, HAND2, MEF2A and SRF conducted in mouse embryonic hearts or cardiac cell types^20,22,50,51^ (see Methods for detail). We found that among the 134 GFP+ specific ATAC-seq peaks that were aCNEs shared with mouse, 49 (37%) are bound by at least one cardiac TF at the orthologous regions of mouse genome and 34 (25%) are CREs that are bound by more than one cardiac TF (Supplementary Table 2). In contrast, cardiac TF binding was rarely observed in GFP- specific open chromatin regions (Supplementary Fig. 3F and Supplementary Table 2).

To assess the *in vivo* function of the GFP+ specific aCNEs, we used a transgenic reporter assay to test their activity during zebrafish embryonic development (Fig. 3C). The regions we selected included both direct (13 regions) and indirect (8 regions) alignments between zebrafish and human (Supplementary Table 3). We cloned the 21 zebrafish regions into a Tol2-based GFP enhancer detection vector and injected the constructs into zebrafish embryos (Fig. 3C).

We found 18/21 regions drove heart expression in at least 30% of the F0 embryos injected (Supplementary Fig. 4), suggesting they are active enhancers in embryonic heart development. Within this set of 18 enhancers, 11 zebrafish aCNEs (ZaCNEs) were located near known cardiac genes, nine of which overlapped the experimentally determined binding sites of one or more cardiac TFs (GATA4, NKX2.5, TBX5, HAND2, MEF2A and SRF) in mouse hearts or cardiac cell types^20,22,50,51^ (Supplementary Table 3). Four of the seven selected ZaCNEs with no known cardiac gene association had experimentally determined binding sites of one or more cardiac TF (Supplementary Table 3).

We raised stable transgenic lines for the 18 ZaCNEs that passed the 30% threshold in the F0 assays, to verify their cardiac activity (Fig. 3C). Except for ZaCNE18 for which only one transgene germline carrier was identified, we obtained multiple independent alleles for the other 17 ZaCNEs (Supplementary Table 3). Despite the random transgene integration mediated by the Tol2 transposase system, we observed consistent GFP expression in hearts in multiple alleles of the same enhancer lines for 15/17 ZaCNEs, with ZaCNE4 and ZaCNE7 being the exceptions (Supplementary Fig. 5). These results from F1 or F2 stable transgenic embryos demonstrated high accordance with those obtained from the F0 generation. We classified the heart expression observed in the ZaCNE transgenic lines into 4 major categories (Fig. 3D and Supplementary Fig. 5A). Some heart enhancers broadly label all heart structures (Category I), while others showed specific or enhanced expression in the ventricle (category II), atrioventricular canal (category III) or outflow tract (category IV) (Fig. 3D and Supplementary Fig. 5). These results confirmed the diverse expression driven by aCNEs and suggested that aCNEs may play distinct roles in regulating heart gene expression.

### Human-zebrafish aCNEs share conserved early cardiac activities

We wanted to determine if human-zebrafish orthologous aCNEs were functionally conserved with respect to their ability to control spatial-temporal gene expression patterns. We chose four aCNEs near essential cardiac genes *hand2/HAND2* (aCNE1), *tbx20/TBX20* (aCNE20) and *mef2cb/MEF2C* (aCNE5, aCNE19). All of these zebrafish sequences drove robust and specific heart expression in stable transgenic lines (ZaCNE1, ZaCNE5, ZaCNE19, ZaCNE20) (Fig. 4A,B and Supplementary Fig. 6A,B). The four orthologous open chromatin regions from human (which have 50-60% sequence identity with their zebrafish counterparts) also drove GFP expression in the hearts of the zebrafish transgenic lines, with 3 of them (HaCNE1, HaCNE5, HaCNE20) demonstrating activities similar to that of their zebrafish orthologs (Fig. 4A,B and Supplementary Fig. 6A). *gfp in*-*situ* characterization further confirmed that the cardiac activities of aCNE1 and aCNE20 human-zebrafish pairs share similar temporal dynamics. Both pairs were active in cardiac lineages at early somite stages (13 hpf, prior to formation of the linear heart tube), 24 hpf, as well as 48 hpf (Fig. 4C,D). Despite marking a broad population within the ALPM (cardiac domain) at 13 hpf, we found that orthologous pairs of enhancers labelled anatomical subregions of the heart in a similar manner at 48 hpf (Fig. 4C,D). The two aCNE1 enhancers both labelled ventricles and the inner curvatures of atria (Fig. 4C-i,i’). For the aCNE20 orthologs, the strongest activity of both was seen in the inner curvature of ventricles and atrioventricular canal regions (Fig. 4D-l,l’).

**Figure 4.**
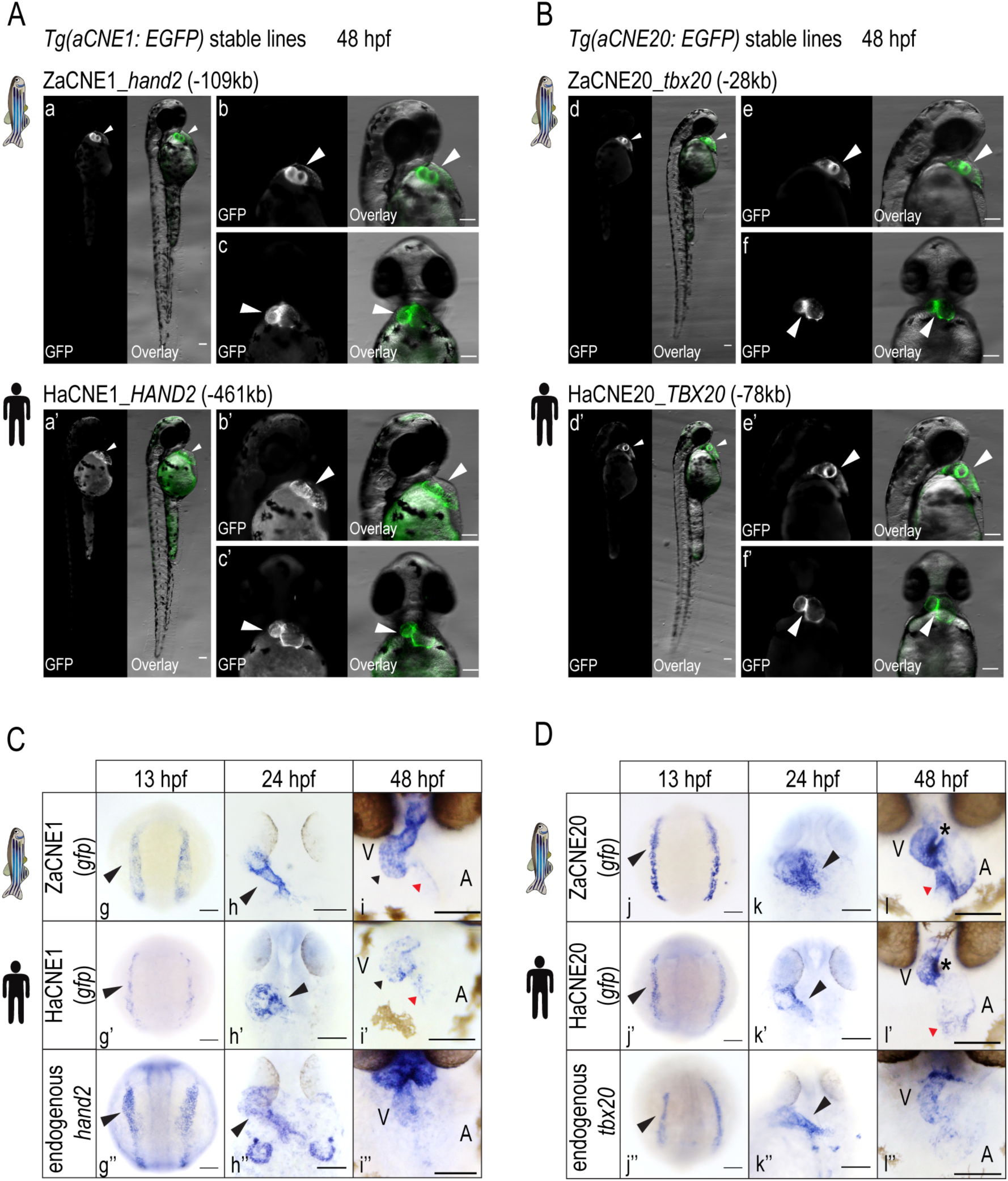
Anciently conserved open chromatin regions share conserved cardiac activities. Fluorescent images (A, B) of aCNE transgenic lines generated using zebrafish (a, b, c, d, e, f) or human (a’, b’, c’, d’, e’, f’) sequences. *In-situ* characterization (C, D) of the activities of the zebrafish (g, h, i, j, k, l) and human (g’, h’, i’, j’, k’, l’) aCNE sequences and the endogenous expression of zebrafish cardiac genes (g’’, h’’, I’’, j’’, k’’, l’’) nearby. In (i) and (i’), black triangles indicate staining in ventricles and red triangles the inner curvature of atria for both aCNE1 transgenic lines. In (l) and (l’), stars indicate the conserved activity of both aCNE20 enhancers at the inner curvature of ventricles and atrioventricular canal regions. Red triangles in (l) and (l’) point to the staining in inflow tract. All images shown were collected from embryos of stable lines and all scale bars represent 100 um.

Our motif enrichment analyses suggested that GATA factors act as important regulators in the GFP+ population. Within GFP+ specific zebrafish aCNEs conserved with human, 66% (107/162) have at least one GATA motif and ~30% (52/162) have more than one GATA motif (Supplementary File. 5). In contrast, while using the same threshold, no significant GATA motifs were found in the conserved GFP- specific zebrafish aCNEs (Supplementary File. 5). To test the functional importance of the GATA motif in aCNEs, we mutated an aligned GATA motif within zebrafish and human aCNE1 regions (near the *hand2/HAND2* locus) and compared their activities with that of the WT sequences (Supplementary Fig. 6C). In F1 stable transgenic lines, we found that both zebrafish and human aCNE1 sequences with mutated GATA motif drove much weaker GFP expression in zebrafish hearts compared to the WT sequences (Supplementary Fig. 6C).

These results together demonstrate that despite sequence divergence between human and zebrafish aCNEs, we have identified a number of conserved accessible chromatin elements that share conserved spatiotemporal, GATA-dependent activity during the early stages of heart development.

### aCNEs are enriched for lineage-specific developmental enhancers

Functional enrichment analysis by GREAT revealed that aCNEs were significantly associated with DNA binding proteins, with the highest enrichment being the homeodomain proteins (GO:0043565, sequence-specific DNA binding: FDR=0; IPR009057, Homeodomain-like: FDR=0; binomial test). Supporting the role of aCNEs as development enhancers, we observed that these regions were also highly enriched for genes regulated by polycomb repressive complex 2 (PRC2) (MSigDB Perturbation, Set ‘Suz12 targets’: FDR=0, Set ‘Eed targets’: FDR=0; binomial test). To gain more insight into lineage-restricted functions of aCNEs we compared the human aCNEs orthologous to our GFP+ specific and GFP- specific zebrafish ATAC-seq peaks. Although human aCNEs specific to the GFP+ and GFP- populations both showed significant enrichments for PRC2 target genes and homeobox transcription factors, only a minority of PRC2 regulated genes were associated with both GFP+ and GFP- aCNEs, indicating that these two sets of aCNEs may be involved in distinct developmental processes (Fig. 5A and Supplementary Fig. 7).

**Figure 5.**
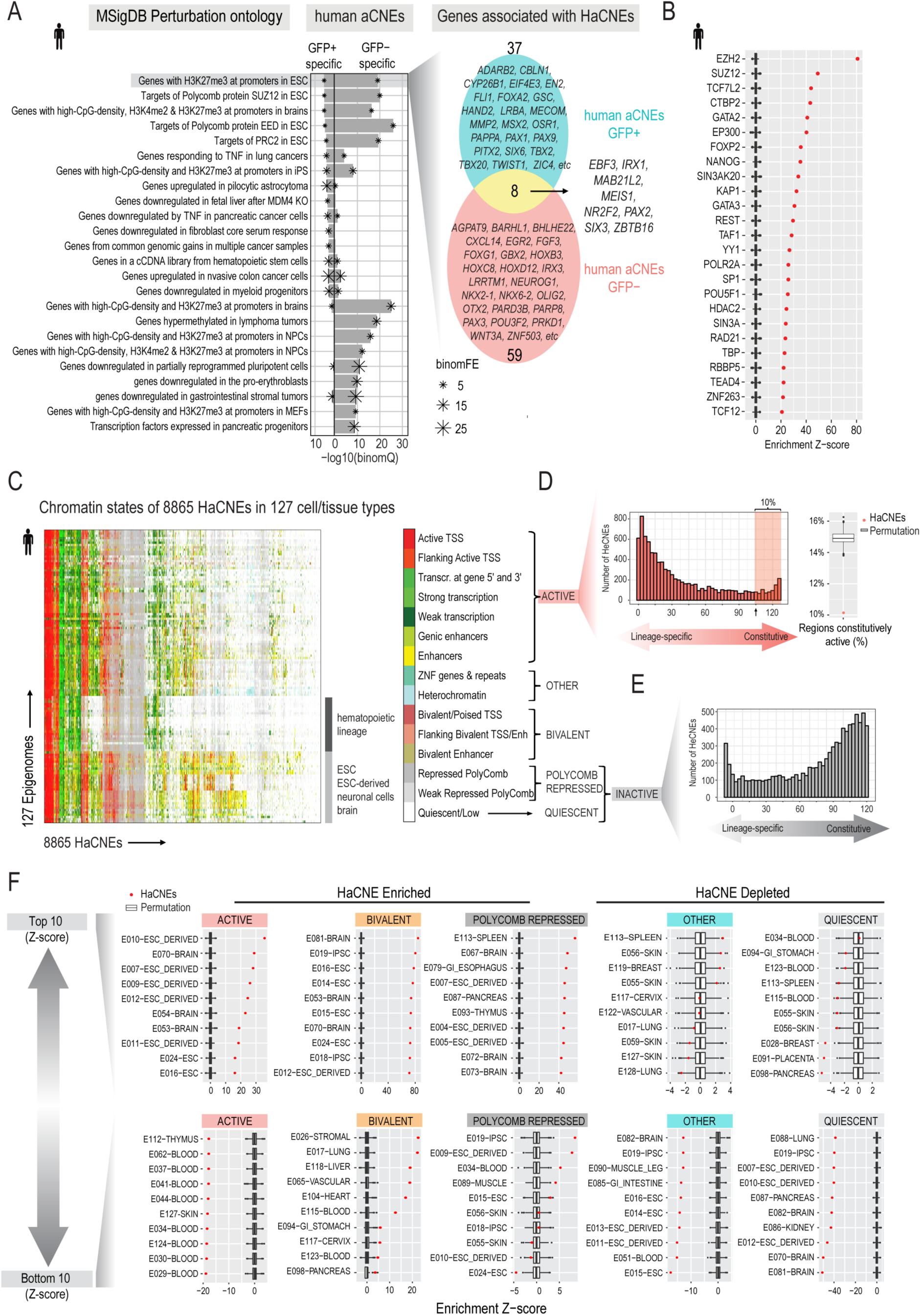
Genomics and epigenomic features of aCNEs. (A) DNA binding factor occupancy enrichment within all HaCNEs (n=8866). Boxplots shows the distribution of 1000 times permutation and red dots represent the enrichment Z-scores normalized by the permutation results (see Methods for details). The top 20 most enriched factors were plotted. (B) Enrichment analysis of HaCNEs conserved with GFP+ or GFP- specific ATAC-seq peaks. Top 15 enrichment terms (binomial FDR < 0.5, FE >2, sorted by binomial FDR) for each category (GFP+, GFP-) were plotted. Both GFP+ and GFP- specific HaCNEs enrich for genes bound by PRC2 subunits and repressed by H3K27me3 modification in ESCs. Venn diagram shows the overlap of genes associated with the GFP+ and GFP- specific HaCNEs that contribute to the top1 enrichment “genes with H3K27me3 at promoters in ESC”. (C) Heatmap showing the chromatin states of 8865 human aCNEs (HaCNEs) across 127 epigenomes from Roadmap Epigenetic Project. One HaCNE on chromosome Y was excluded from this analysis since it was absent in several epigenomes from female tissues, which left 8865 regions in this analysis. Each column represents a HaCNE and each row an epigenome. (D) Histogram shows the distribution of HaCNEs that displayed ACTIVE chromatin states (x-axis). Approximately 10% of HaCNEs were constitutively active in the majority of the epigenomes (>80%) (shaded area in histogram). Compared to the same number of randomly selected open chromatin regions from the ENCODE3 human DHS master list, HaCNEs were depleted for regions constitutively active (red dot on boxplot on the right). (E) Histogram of HaCNEs that displayed QUIESCIENT or POLYCOMB REPRESSED states. (F) Analyzing if certain chromatin states (ACTIVE, BIVALENT, POLYCOMB REPRESSED, QUIESCENT and OTHERS) were enriched in HaCNEs compared to randomly selected open regions (permutation). The enrichment Z-score were plotted for each epigenome (red dot) along with the permutation distribution (boxplot) (see Methods for details). For each chromatin state, epigenomes with the top 10 (up panel, most enriched/least depleted) and bottom 10 (down panel, most depleted/least enriched) Z-score were plotted. Row names were formatted as “ID”-”EpigenomeName”, which were retrieved from Roadmap Epigenomics Project metadata table (http://egg2.wustl.edu/roadmap/web_portal/meta.html).

To characterize TF occupancy preferences within aCNEs, we used the human ENCODE TF binding sites (TFBS) dataset which contains uniformly analyzed ChIP-seq binding profiles for 161 factors in 91 cell types (Transcription Factor ChIP-seq Clusters V3 from ENCODE). We asked if any factor showed occupancy enrichment within aCNEs compared to randomly selected open chromatin regions (see Methods for details). The factors with the highest enrichment Z-scores were subunits of PRC2 (EZH2, SUZ12) (Fig. 5B). Other transcription repressors (CTBP2, SIN3A, REST, KAP-1) or dual regulators (TCF7L2, YY1, associated with both active and repressive regulation in different contexts) were also seen among the top 20 most enriched factors (Fig. 5B). We also observed significant enrichments for a wide-variety of transcription factors (GATA2, FOXP2, NANOG) and factors controlling chromatin architectures (RAD21, CTCF) (Fig. 5B and Supplementary Fig. 7), suggesting aCNEs play diverse roles in gene regulation.

To get a global understanding of the tissue-specific usage of aCNEs, we took advantage of the chromatin states of 127 human tissues/cell types that were predicated based on 5 chromatin marks (H3K4me3, H3K4me1, H3K36me3, H3K27me3 and H3K9me3) in the Roadmap Epigenomics Project^52^ (Fig. 5C and Supplementary Fig. 8). We simplified the 15 chromatin states into “Active”, “Bivalent”, “Polycomb repressed”, “Quiescent” and “Other” categories (Fig. 5C and Supplementary Fig. 8). Only 10% (899/8865) of the aCNEs showed constitutive activity (“Active” in > 80% the tissues/cell types) (Fig. 5D). Supporting this observation, aCNEs were depleted for regions of constitutive activity compared to bulk open chromatin regions (ENCODE3 human DNase I hypersensitivity site Master list, *P* < 0.001, permutation test; Fig. 5D). Supporting their function as lineage-restricted enhancers, we found more than half of aCNEs to be “Active” in a lineage specific manner (active chromatin state in at least one epigenenome and Polycomb repressed or Quiescent in > 70% of the epigenomes (4603/8865), Fig. 5D,E). To examine if certain chromatin states were enriched in aCNEs, we compared the percentage of a chromatin state among aCNEs to that among randomly selected open chromatin regions in each epigenome. We found that aCNEs were enriched for “Bivalent” (126/127 epigenomes) and “Polycomb repressed” (121/127 epigenomes) chromatin and depleted for “Quiescent” (120/127 epigenomes) and “Other” (119/127 epigenomes) chromatin regions in the vast majority of the epigenomes (Fig. 5F and Supplementary Fig. 8C). For “Active” chromatin regions, aCNEs showed enrichment in some epigenomes (brain, neuronal cell types) but depletion in more tissue/cell types (hematopioetic lineages) (Fig. 5F and Supplementary Fig. 8C).

Given that lineage-specific activity and enrichment for PRC2 binding are well-established properties of poised enhancers in ESCs^53–56^, we compared aCNEs to three enhancer types defined in a recent study using mouse ESCs: poised (P300+, H3K27me3+, H3K4me3-, H3K27ac-), primed (H3K4me1+, H3K4me3-, H3K27ac-, H3K27me3-) and active (P300+, H3K27ac+, H3K4me3-, H3K27me3-)^56^. Compared to all open chromatin regions, aCNEs showed a 5-fold enrichment for poised enhancers (FDR < 4.44e-19 using hypergeometric test) with little enrichment for active (1.4 fold FDR < 0.0018 using hypergeometric test) or primed (1.1 fold FDR < 0.16 using hypergeometric test) enhancers. We also observed a strong enrichment for poised enhancers that were identified in human ES cells (fold of enrichment: 9.72, FDR < 1.14e-84 using hypergeometric test)^54^.

Conserved genomic regulatory blocks (GRBs), which are defined by cluster of CNEs, contain developmental genes, coincide the boundaries of topologically associating domains (TADs) and remain concordant between species of various evolutionary distances^57^. We found that nearly 90% (7846/8866) of human aCNEs fell into GRBs defined by using CNEs (70% identity over 50bp) between human and chicken^57^, suggesting aCNEs have been kept in a discrete set of syntenic regions during vertebrate evolution. Overall the genomic and epigenomic properties of aCNEs suggest that many of them may serve as lineage-restricted enhancers that facilitate the expression of developmental genes.

## Discussion

The existence of a phylotypic period implies that the gene regulatory events that control phylotypic gene expression patterns are also likely to be evolutionarily constrained. In this study we characterized a population of cells enriched for cardiac progenitors in early zebrafish embryos. By studying the open chromatin regions in this population, we uncovered a set of putative enhancers. Leveraging two resources that characterized non-coding elements conserved between zebrafish and humans^25,26^ allowed us to identify more than 6000 zebrafish open chromatin regions that overlapped open chromatin regions in human or mouse genomes. Of these conserved accessible chromatin regions (aCNEs), 162 were unique to our cardiac progenitor-cell enriched population. While 27% (43/162) of the orthologous human regions have recently been predicted to be cardiac enhancers ^23^, to our knowledge none of these 162 zebrafish open chromatin regions or their orthologous human or mouse sequences had been previously tested *in vivo*. We found that 15/21 these pre-phylotypic aCNEs drove cardiac expression in stable zebrafish lines at 24hpf, which is around the time considered to be the zebrafish phylotypic stage3,4,7 or later in development (48hpf). Overall our data support the existence of conserved cis regulatory elements that are primed early in development prior to the establishment of the body plan and function during the phylotypic stage.

Many functional CREs do not share overt sequence similarities due to rapid turnover of the spacing between, and sequence of TF binding motifs^45, 58–61^. This phenomenon is prevalent even within rodent^62^ or primate^47^ orders, let alone comparisons between species separated over large evolutionary distances, such as zebrafish and human (over 400 million years). Nonetheless, sequence comparisons between human and teleosts over great phylogenetic distance have discovered many functional non-coding elements active during development^26, 63–66^ and CNEs are often found near genes encoding developmental regulators^25,26,63,65,67,68^. More direct evidence of conserved cis regulatory events acting prior to the phylotypic period have come from profiling epigenetic changes around the phylotypic period. For example, zebrafish embryos have shown that regions dynamically gaining epigenetic modification indicative of development enhancers (H3K27ac) at gastrulation stage (8.5hpf) are enriched for evolutionarily conserved DNA sequences^6^. Furthermore, differential DNA methylation changes observed during the phylotypic period in zebrafish, mouse, and frog were enriched for evolutionarily constrained DNA sequences and these DNA methylation changes were presumably guided by prior sequence-specific transcription binding events^7,69^. Our results identify a substantial number of aCNEs that are established in the early embryo, many of which drive tissue specific gene expression patterns later in development.

Vertebrate heart enhancers tend to show a lack of overt phylogenetic conservation relative to enhancers active in other tissues, such as the brain, at the same developmental stages^17,41^. Our results echo this finding, while at the same time highlighting a set of highly conserved heart enhancers. By comparing open chromatin within CNEs identified using approaches such as transitivity through a third species and ancestry reconstruction^25,26^, we discovered 60% more conserved open chromatin regions between zebrafish and human/mouse that would otherwise have been missed by using direct sequence alignment alone (Fig. 3A,B). Nearly half of the heart enhancers we validated were from indirect sequence alignment (Supplementary Table 3), and zebrafish-human aCNE orthologs identified from either direct (aCNE1) or indirect (aCNE5, aCNE19, aCNE20) alignment share conserved cardiac activity to a similar degree, despite a 50-60% sequence identity. The aCNEs that we have validated can be used to further our understanding of heart development and regeneration. For example, using one of these “indirectly aligned” heart enhancers (aCNE21) we can recapitulate the endogenous expression of the nearby cardiac gene *hey2*^70^. Characterizing the epigenomic landscape of additional cell-lineages early in development will likely highlight further enhancers, and reveal specific aCNEs capable of driving cell or tissue-specific expression before, during and after the phylotypic period.

Attesting to their potential importance in regulating specific genes through long-range chromatin interactions, an established genomic property of CNEs is that they cluster into regions of conserved synteny, referred to as gene regulatory blocks (GRBs)^71,72^. A significant fraction of GRB boundaries coincide with the boundaries of topological association domains (TADs)^57^. We found that nearly 90% (7846/8866) of our human aCNEs fell into GRBs comprised of CNEs between human and chicken (70% identity over 50bp)^57^. Overall the genomic and epigenomic properties of aCNEs suggest that many of them may serve as lineage-restricted enhancers that facilitate the expression of developmental genes.

We conclude that conserved open chromatin regions established prior to the phylotypic period, and shared over 450 million years of evolution, likely represent a set of ancient enhancers that contribute to diverse spatial and temporal gene expression patterns. Although the first deletions of ultraconserved non-coding elements did not reveal overt phenotypes^73^, new studies are beginning to demonstrate developmental anomalies^74,75^. Consistent with the existence of shadow enhancers, which can buffer the effects of individual enhancer loss^76–79^, two recent studies showed that the pairwise deletion of ultraconserved enhancers had an increased phenotypic impact^75,79^. While more work remains to be done to dissect the *in vivo*, spatial-temporal expression patterns and function of anciently conserved vertebrate enhancer, regions of deeply conserved open chromatin represent a solid foundation from which to explore the regulation and evolution of the vertebrate body plan.

## Methods

### Zebrafish husbandry and line maintenance

Zebrafish were maintained and handled under the guidance and approval of the Canadian Council on Animal Care and the Hospital for Sick Children Laboratory Animal Services. Embryos were raised at 28.5 degree Celsius and staged as previously described^80^.

### Zebrafish injection and transgenic lines

2.7 kb Mouse *Smarcd3*-F6 sequence (mm9 assembly chr5: 24113559-24116342)^13^ was sub-cloned from a gateway entry vector into Zebrafish Enhancer Detection (ZED) Vector^81^ and *E1b-Tol2-GFP-gw* vector^82^ (Addgene #37846) using Gateway recombination system (Invitrogen, Gateway LR Clonase II Enzyme Mix, Cat# 11791020). Tol2 mRNA was synthesized using mMessage mMachine kit (Ambion, Cat# 12538120) and purified by MEGAclear™ Transcription Clean-Up Kit (Ambion, Cat# AM1908). 25ng ZED or *E1b-Tol2-GFP-gw* plasmid carrying the *Smarcd3*-F6 enhancer was microinjected into wildtype embryos at one-cell stage with 150 ng Tol2 mRNA following the standard Tol2 procedure^83^. 2 independent germline carriers have been identified for *Tg (Smarcd3-F6: EGFP*) generated using *ZED-Smarcd3-F6* plasmid and 4 independent carriers have been identified from F0 founders generated using *E1b-Tol2-GFP-Smarcd3-F6* plasmid. Transgenic embryos from different carriers display similar GFP expression in heart. 4 alleles (*Smarcd3-F6:EGF^Phsc70^, Smarcd3-F6:EGF^Phsc71^, Smarcd3-F6:EGF^Phsc72^, Smarcd3-F6:EGF^Phsc73^*) have been raised (see Supplementary Table 3 for details). A line (*Smarcd3-F6:EGF^Phsc70^*) with brighter GFP expression (generated using ZED-Smarcd3-F6 plasmid) was used for all related figures presented.

To generate a *Tg(Smarcd3-F6:CreERT2*) line, we adopted two strategies. In the first one, EGFP sequence in *E1b-Tol2-GFP-gw* vector was first replaced by CreERT2 sequence to generate an *E1b-Tol2-CreERT2-gw* vector. Then *Smarcd3*-F6 enhancer was sub-cloned into this CreERT2 vector via Gateway recombination. In the second strategy, *Smarcd3*-F6 enhancer was sub-cloned into a gateway 5’ entry vector (*p5E-Smarcd3-F6*). A middle entry vector containing the gata2 minimal promoter that was used in the ZED vector (*pME-gata2P*) and a 3’ entry vector with CreERT2 (*p3E-CreERT2*) were also generated. *p5E-Smarcd3-F6, pME-gata2P* and *p3E-CreERT2* were recombined with *pDestTol2-cCrystalGFP* to generate *pDestTol2cCrystalGFP:Smarcd3-F6-gata2P-CreETR2* construct (Invitrogen, LR Clonase II Plus Enzyme, Cat# 12538120). All gateway entry vectors used were from the Tol2 kit^84^. Microinjection was carried out in the same way as for *Tg (Smarcd3-F6: EGFP*) line. 5 independent lines from *E1bTol2CreERT2:Smarcd3-F6* and 2 independent carriers from *pDestTol2cCrystalGFP:Smarcd3-F6-gata2P-CreETR2* were identified. Eventually 5 different alleles (*Smarcd3-F6:CreERT2^hsc74^, Smarcd3-F6:CreERT2^hsc75^, Smarcd3-F6:CreERT2^hsc76^, Smarcd3-F6:CreERT2^hsc81^, Smarcd3-F6:CreERT2^hsc88^*) were raised (see Supplementary Table 3 for details). Lineage tracing results seen in different *Tg(Smarcd3-F6:CreERT2*) lines were not obviously different. Results presented in Supplementary Fig. 1 were generated from embryos of *Tg (Smarcd3-F6:CreERT2)^hsc76^*. *Tg(nkx2.5: ZsYellow)^fb7^* line used have been previously described^85^.

### RNA probe synthesis and in-situ hybridization

EGFP sequence from the ZED vector and Cre sequence from *p3E-CreERT2* were sub-cloned into pGEM Teasy vector (Promega, Cat# A1360) to make antisense probes (EGFP-F: GGATCCATGGTGAGCAAGGGCGAGGAG, EGFP-R: CTCGAGTTACTTGTACAGCTCGTCCATGCCG; Cre-F: GGCGTTTTCTGAGCATACCTG, Cre-R: CCCAGGCTAAGTGCCTTCTCT). DIG-labeled in-situ probes were synthesized using DIG RNA Labeling kits (Roche). RNA in-situ was carried out as previously described ^86^. Stained embryos were cleared in BBA solution (2:1 Benzyl benzoate: Benzyl alcohol) and imaged under a Leica M205FA stereomicroscope

### Immunostaining

Immunostaining was carried out as previously described with minor modifications^87^. Embryos from *Tg (Smarcd3-F6: EGFP)^hsc70^* and *Tg(nkx2.5: ZsYellow)^fb7^* crosses were fixed at 13 hpf in 4% paraformaldehyde at 4 degrees Celsius overnight. After 3X5min washes in PBS with 0.1% Triton, embryos were permeabilized in PBS with 0.5% Triton for 4 hours at room temperature (RT). Embryos were then blocked in PBST (1% DMSO and 0.5% Triton X in PBS) with 5% Normal Goat Serum (Millipore, Cat# S26-LITER) for 1.5-2h at RT before incubated with primary antibodies (α-RCFP 1:500, Clontech Cat# 632475; α-GFP 1:1000, Torrey Pines Biolabs) at 4 degrees Celsius overnight. Next day, embryos were washed for 3-4 hours in PBS with 0.1% Triton at RT, with 6-8 changes of solution. Incubation of secondary antibodies was carried out at 4 degrees Celsius overnight, followed by the same washing procedure as that for primary antibodies. After staining, embryos were mounted in 1% (w/v) low melt agarose (Sigma, A9414) and imaged under a Nikon A1R Si Point Scanning Confocal.

### CreETR2 lineage tracing experiment

Embryos from *Tg (Smarcd3-F6:CreERT2)^hsc76^* and *Tg(βactin2:loxP-DsRed-STOP-loxP-EGFP)^s928Tg^* crosses were dechorionated and incubated in 5uM 4-OH-Tamoxifen(4-HT, Sigma cat# T176) for 12 hours. The time when 4-HT was added was specified in Supplementary Fig. 1. After treatment, embryos were rinsed twice, placed in fresh egg water and imaged at 48 hpf under Zeiss Axio Zoom.V16 Stereoscope.

### Embryo dissociation, Fluorescence activated cell sorting (FACS)

Around 100 *Tg(Smarcd3-F6: EGFP)^hsc70^* embryos were dechorionated with pronase (Sigma, Cat# 11459643001) and transferred to a 1.5mL Eppendorf tube when they reached bud stage. After incubated in 200ul calcium-free Ringer solution (116 mM NaCl, 2.6mM KCl, 5mM HEPE, pH 7.0) for 5 min, embryos were transferred into a 24-well plate filled with 500 ul TrypLE solution (GIBOCO, TrypLE Express Enzyme, cat #: 12604-013) for dissociation at room temperature. Embryos were gently homogenized every 5min with P1000 tips. Dissociation was monitored under a dissection scope until most cells were in a single-cell suspension. The cell suspension was transferred into 200ul ice-cold fetal bovine serum (FBS) to stop the reaction. Cells were centrifuged at 300g for 3min at 4 degree and washed with 500 ul ice-cold DMEM with 10% FBS before resuspended in 500 ul ice-cold DMEM with 1% FBS. Right before the FACS, cells were filtered through a 40um strainer and DAPI was added at a concentration of 5 ug/ml to exclude dead cells.

FACS was performed on Beckman Coulter Mo Flo XDP or Mo Flo Astrios sorter with a 100um nozzle by the SickKids-UHN Flow and Mass Cytometry Facility. Cell-doublets and dead cells were excluded based on forward scatter, side scatter and DAPI channel. GFP+ and GFP- cells were sorted into 100% FBS and subjected to RNA-seq or ATAC-seq procedures immediately once sorting finished. 30,000-50,000 GFP+ cells and 100, 000 GFP- cells were usually collected in one sort.

### Bulk mRNA-seq and single-cell mRNA-seq

Single-cell cDNA libraries were prepared using Fluidigm C1 system. After FACS, GFP positive cells were washed twice in DMEM with 3%FBS and filtered through a 40um cell strainer. Cells were adjusted to a concentration of 400-500 cells/ul before mixed with C1 suspension solution at a 5.2:4.8 ratio. Then 10ul final cell mixture was loaded into a C1 medium or small Chip. Cell capture was examined under a microscope and only wells with a single cell captured were included in library construction. 3 ArrayControl™ RNA Spikes were added into the cell lysis mixture according to Fluidigm C1 single-cell mRNA-seq protocol (PN 100-7168 Rev. B1). Cell lysis, Oligo-dT primer mediated reverse transcription, 21 cycles of PCR amplification and cDNA harvesting were performed as per manufacture’s instruction (Fluidigm, PN 100-7168 Rev. B1). We usually recovered 30-40 cells (30-40% capture efficiency) from one C1 Chip, eventually 96 single-cell cDNA libraries were collected from three batches of experiments at the end.

For bulk mRNA-seq, RNA from 4000 GFP+ or GFP- cells were prepared using RNeasy Plus Micro Kit (Qiagen, Cat# 74034) and converted into cDNA libraries following the Fluidigm tube control protocol (the same protocol as that for single-cell mRNA-seq except the input cell number is different). Three biological replicates were collected for both GFP+ and GFP- samples.

For both single-cell and bulk mRNA-seq, final sequencing libraries were made using Nextera XT DNA Sample Preparation Kit (Illumina, Cat# FC-131-1096) and 120bp pair-end sequenced on a Illumina HiSeq 2500 platform according to manufacturer’s instruction. Bulk RNA-seq libraries were sequenced to a depth of (18±1.9) million reads and single-cell mRNA-seq libraries a depth of (3.0±0.7) million reads.

### ATAC-seq

ATAC-seq was performed following published protocol^88^ with minor modifications. 30,000-50,000 cells obtained from FACS were used for nuclei prep. After tagmentation reaction, transposed DNA fragments were amplified using the following PCR condition (1 cycle of 72 °C for 5 min and 98 °C for 30 s, followed by 12 cycles of 98 °C for 10 s, 63 °C for 30 s and 72 °C for 1 min). Amplified libraries were purified twice with Agencourt Ampure XP beads (beckman coulter, Cat#A63880) with a bead-to-sample ratio of 1.8:1. Qubit fluorometer and Aglient Bioanalyzer were used to check library quality and concentration. Libraries were 50bp single-end sequenced on Illumina HiSeq 2500 platform to a depth of (3.5-7.0) × 10^7^ reads. Two biological replicates were collected for both GFP+ and GFP- cells.

### Transgenic zebrafish enhancer assay

Candidate regions containing the zebrafish ATAC-seq peaks (21 ZaCNEs) or human DHSs (4 HaCNEs) were amplified from genomic DNA and recombined into *pDONOR221* vector (Invitrogen, Gateway BP Clonase II Enzyme Mix, Cat# 11789020) before they were cloned into *E1b*-*Tol2*-*GFP*-*gw* vector eventually. 25 ng *E1b*-*Tol2*-*GFP*-*gw* plasmid carrying one aCNE and 150ng Tol2 mRNA were injected into WT embryos at one-cell stage. F0 founder embryos were raised to 48-52 hpf before subjected to imaging and heart expression scoring. Candidate regions were considered as a heart enhancer if more than 30% of the injected embryos display GFP positive cells beating in zebrafish hearts, which was consistent with the criterion used in similar studies before^89^. 42-220 (average n=86) injected embryos were analyzed for each candidate regions. The genomic coordinates, lengths and nearby genes of candidate regions and primers used for cloning can be seen in the Supplementary Table 3.

F0 embryos injected with enhancers that passed the 30% threshold were raised to screen for transgene germline carriers. Except enhancer ZaCNE18 for which only one carrier has been identified, 2-4 independent alleles have been identified for each enhancer (see Supplementary Table 3). Though ectopic expression has been seen in some carriers, the cardiac expression patterns were similar in different alleles of the same transgene.

### GATA motif mutagenesis

GATA motif mutation was introduced by primers designed by Agilent Genomics QuickChange program (http://www.genomics.agilent.com/primerDesignProgram.jsp) (ZaCNE1_muta_F: cagattaggacccagctaggtgccagtggggggggtgttagtgcagaaaaggttacactac; ZaCNE1_muta_R: gtagtgtaaccttttctgcactaacaccccccccactggcacctagctgggtcctaatctg; HaCNE1_muta_F: attagagtgaaaagaggtgccggtggggggggtgcgaatgcgccaggggtcacgc;

HaCNE1_muta_R: gcgtgacccctggcgcattcgcaccccccccaccggcacctcttttcactctaat).

The primers were designed to convert the aligned GATA consensus sequence AGATAA to CCCCCC. *pDONOR221* vectors carrying the mutated aCNE1 enhancers were PCR amplified from the original *pDONOR221* containing the wildtype (WT) aCNE1 sequences using the primers above. After amplification, 50 ul PCR mixture was incubated with 1 ul DpnI (NEB, Cat#R0176S) at 37 degrees to remove WT enhancer templates. DpnI was inactivated at 80 degrees for 20min before transformation. Plasmid clones with the correct GATA motif mutation were confirmed by Sanger sequencing and used as entry clones to make *E1b-Tol2-GFP* constructs containing mutated aCNE1 enhancers. Independent germline carriers haven been identified for ZaCNE1_GATAMutated (n=5) and HaCNE1_GATAMutated (n=4) enhancers (see Supplementary Table 3). GFP expression levels in different GATA_Mutated alleles look slightly different but were generally weaker than WT_alleles. Images in Supplementary Fig. 6 were taken using alleles showing median expression level within all alleles.

### Processing and analysis of bulk and single-cell mRNA-seq data

Raw reads were analyzed with FastQC (version 0.11.2)^90^, trimmed with Trimmomatic (version 0.32)^91^ (ILLUMINACLIP:TruSeq3-PE-2.fa:2:30:10:5:true LEADING:20 TRAILING:20 SLIDINGWINDOW:5:25 MINLEN:36) before aligned to Zv9 zebrafish genome assembly using STAR^92^ using default setting. The percent of uniquely mapped reads is (80±2.6)% for bulk samples and (63±9.3)% for single-cell libraries (see Supplementary Table 4 for details). HTSeq-count^93^ and Zv9 (Ensembl release 79) transcriptome annotation were used to determine the number of reads mapped to each gene. An average of 14486 genes were detected in bulk samples and 3901 genes in single cells.

For bulk mRNA-seq, genes that have at least 1 read per million in at least 2 replicates were kept for downstream analysis. edgeR package (version 3.18.1) was used for normalization and differential gene expression analysis^94^. 167 genes were identified as more highly expressed in GFP+ cells compared to the GFP- population while 147 genes were more strongly expressed in GFP- cells (FDR < 0.05, Fold change > 2). Volcano plot showing differentially expressed genes was generated with an in-house R script.

Functional enrichment analysis was performed using online tool g:Profiler^95^. A more stringent list of differentially expressed genes (FDR < 0.05, Fold change > 4) were ordered based on its fold change and then used as the input for g:Profiler. Only functional categories containing more than 2 genes but less than 500 genes were included in our analysis. Benjamini-Hochberg FDR method was used for multiple testing correction to adjust significance thresholds. The top10 enriched GO terms (sorted by FDR) for each gene list were plotted.

For single-cell data, 92 cells with more than 2000 genes detected (counts per million, CPM >0) were kept for clustering analysis. Gene counts (CPM) for each cell were first normalized using TMM method^96^ and then winsorized before log transformation (In(CPM+1)). To perform unsupervised clustering, 189 genes were selected and used, if they were differentially expressed in bulk mRNA-seq (|log2FC|>1.5, FDR < 0.1) and have an expression level (In(CPM+1) > 4) in at least one single cells. The Manhattan distance and ward.D method were used for unsupervised clustering. To perform PCA analysis on single cells, R package FactoMineR^97^ was used with default settings.

### ATAC-seq reads mapping, peak calling and differential peak identification

Raw reads were preprocessed by FastQC^90^ and Trimmomatic^91^ (ILLUMINACLIP:/hpf/tools/centos6/trimmomatic/0.32/adapters/TruSeq3-PE-2.fa:2:30:10:5:true LEADING:20 TRAILING:20 SLIDINGWINDOW:5:25 MINLEN:36) before aligned to Zv9 zebrafish genome assembly by BWA (version 0.7.8) under aln model^98^. Reads with mapping quality score > 30 were kept for downstream analysis (samtools view -b -q 30). Two replicates for each sample were merged for peak calling by MACS2^99,100^ (version 2.7.9) (--nomodel -- nolambda --gsize 1.4e9). Around 150,000 peaks were identified in both GFP+ and GFP- populations. Peaks showing enriched signals in one population versus the other (GFP+/- specific peaks) were identified using DiffBind package^101^ (DEseq2, FDR < 0.05). After DiffBind analysis, a total of 200,937 ATAC-seq peaks have been identified, within which 3838 are GFP^+^ specific peaks and 1633 are GFP^-^ specific ones.

### Motif enrichment and functional enrichment analysis

Motif enrichment analysis was conducted using CentriMO^102^ (version 4.11.2). In order to obtain the summits of GFP+/- specific peaks, we first perform BEDtool-intersect between the peak summits determined by MACS2^99,100^ and the GFP+/- specific peaks identified by diffBind^101^. By doing this, we identified 3861 GFP+ specific summits and 1635 GFP- specific ones. 250bp was extended to each side from the peak summits and the new regions with 501 bp uniform length were used as input for CentriMO. To identify motifs enriched in GFP+(-) specific peaks, the same numbers of shared peaks were subsampled and used as the comparative dataset in the “absolute and differential model”. All other parameters were kept as default. Fisher E-values of the top 4 enriched motifs were plotted in the motif probability graphs.

GREAT (version 3.0.0) online tool^30^ was used for functional enrichment analysis of open chromatin regions. Whole genome was used as background and “basal plus extension” rule was used for associating genomic regions with genes. Functional terms showing a binomial FDR < 0.05 and a minimal region-based fold of enrichment of 2 were ranked by their FDR and selected for plotting.

### Identification of sequence-conserved open regions

Two zebrafish CNE datasets were used for this analysis. First, Zebrafish CNEs (zCNEs) conserved with human or mouse identified from Hiller et al. were obtained from the authors, including records of direct sequence alignment and transitive mapping through other species or reconstructed ancestries^25^. zCNEs that overlaps a zebrafish-human well-aligning window^25^ by at least 15bp were defined as direct aligned. zCNEs that can be mapped back to mouse through transitive alignment or ancestry reconstruction but cannot be detected by direct alignment were defined as indirectly aligned CNEs. The same direct and indirect alignment definition was set for zCNE conserved with mouse (zCNE_mouse)

To include more high-quality zebrafish CNEs into our analysis, we used another zebrafish-human CNE dataset identified in a recent study through transitive alignment via the spotted gar genome^26^, which we referred to as “garCNE”. If the zebrafish coordinate of a garCNE does not overlap any zCNE records, we added it into our analysis and define it as indirect aligned. Altogether, we collect 20,005 zebrafish CNEs conserved with human, with 10187 direct and 9818 indirect ones (Supplementary Table 1).

To establish the same system for CNEs conserved between zebrafish and mouse, we used liftOver to convert the human-zebrafish CNEs identified through gar genome to mouse-zebrafish CNEs (-minMatch=0.1), and then add those CNEs on top of zCNE_mouse dataset in a similar way as what we did for human. At the end we obtained 18,827 zebrafish CNEs conserved with mouse, with 9429 direct and 9399 indirect ones (Supplementary Table 1). Finally, we associated zebrafish ATAC-seq peaks to CNEs (sequence-conserved ATAC-seq peaks) if they overlap a CNE by at least one base pair. Nearly 90% of the these CNE-associated ATAC-seq peaks were completely excluded from gene coding regions defined by Ensembl transcriptome annotation (Zv9, release 79).

### Identification of aCNEs

Human or mouse DNase master peak lists from ENCODE3^48^ (https://www.encodeproject.org/search/?type=Annotation&annotation_type=DNase+master+peaks) were used for identifying open regions anciently conserved between zebrafish and human or between zebrafish and mouse. Mouse DNase master peaks, which were provided as mm10 coordinates, were converted to mm9 coordinates using liftOver (-minMatch=0.95). If a CNE that overlaps a zebrafish ATAC-seq peak by at least one basepair also overlaps a DNase I hypersensitivity site (DHS) from the DNase master lists by at least one basepair, then these orthologous ATAC-seq peak and DHS linked via CNEs were identified as accessible CNEs (aCNEs).

There are 1,199,722 non-overlapping regions in mouse DHSs master list after liftOver, 4,193,929 non-overlapping regions in human DHS master list. Probably due to the total number differences, more human DHSs have been identified as aCNEs than mouse DHSs; similarly, more ATAC-seq peaks were identified as aCNEs shared with human than those with mouse (Supplementary Table 1).

We found analysis with the DHS master lists sometimes gave us a chain of DHSs that are conserved with the same ATAC-seq peaks. To avoid potential bias that may be introduced in analysis conducted by GRAET, we merged the DHSs that are within 200bp distance and conserved with the same ATAC-seq peaks. We used merged DHSs for analyses in Fig. 5A, Supplementary Fig. 3F, and Supplementary Fig. 7B,C. All other analyses were conducted on original DHS coordinates.

### Annotating aCNEs with chromatin states of 127 epigenomes

Chromatin states that were defined by 5 chromatin marks (H3K4me3, H3K4me1, H3K36me3, H3K27me3, H3K9me3) for 127 human tissuess/cell types were retrieved from Roadmap Epigenomics Mapping Consortium (http://egg2.wustl.edu/roadmap/web_portal/chr_state_learning.html). Human aCNEs were intersected with the dense bed files for each of the 127 epigenomes (http://egg2.wustl.edu/roadmap/data/byFileType/chromhmmSegmentations/ChmmModels/coreMarks/jointModel/final/) using BEDtools^103^ (intersectBed, Version 2.19.1) to determine the chromatin state of each aCNE region in different epigenomes. If one aCNE region overlaps multiple chromatin states in one epigenome, the state that shared the longest overlap with the aCNE were attributed to the aCNE. Each chromatin state was represented by an integer ranging from 1 to 15, which was defined by Roadmap Epigenomics Project. Clustering was performed with Manhattan distance and Ward.D clustering methods in R. Heatmap showing the chromatin state of the each human aCNE region were plotted using the same color scheme as that used by Roadmap Epigenomics Project.

To simply the analysis, we collapsed the 15 chromatin states into 5 major categories, including “Active” (1: Active TSS, 2: Flanking Active TSS, 3: Transcr. at gene 5’ and 3, 4: Strong transcription, 5: Weak transcription, 6: Genic enhancers, 7: Enhancers), “Bivalent” (10: Bivalent/Poised TSS, 11: Flanking Bivalent TSS/Enh, 12: Bivalent Enhancer), “Polycomb repressed” (13: Repressed Polycomb, 14: Weak Repressed PolyComb), “Quiescent” (15: Quiescent/Low) and “Others” (8: ZNF genes & repeats, 9: Heterochromatin). If an aCNE region was associated with an “Active” state in more than 80% of the 127 tissues/cell types, the aCNE was considered as constitutively “Active”. If an aCNE region were “Polycomb repressed” or “Quiescent” in more than 70% of the 127 tissues/cell types and “Active” in at least one epigenome, it was considered as “Active” in a lineage specific manner.

### Assessing the genomic and epigenomic features of anciently conserved open regions via permutation analysis

Details of permutation design for anciently conserved open regions in each species can be found in this table below.

|  | aCNEs | Permutation |
| --- | --- | --- |
| ATAC-seq peaks | 4496 ATAC-seq peaks that were identified as aCNEs conserved with both human and mouse | 4496 regions randomly selected from all ATAC-seq peaks for each permutation |
| Human ENCODE3 | 8866 DHSs that were identified as aCNEs | 8866 regions randomly selected from human DNase master list |
| DNase sites | conserved with zebrafish | for each permutation |

We randomly subsampled the same of number regions from the whole ATAC-seq peak set or DHS sets for 1000 times and then compare their genomic features (GC content, region length, TF binding occupancy, chromatin states etc.) with that of aCNEs. Average GC contents were calculated using BEDtools^103^ (nucBed, Version 2.19.1).

For TF occupancy comparison, the TFBS cluster data from ENCODE was used, which contains the binding sites of 161 human TFs/co-factors in 91 different cell types determined by ChIP-seq experiments. If a region overlaps a record from TFBS cluster data by at least one basepair, it is counted as evidence of binding event at that region. We counted the total number of binding events exists in each set of regions (aCNEs VS random selected) for each factors included in TFBS cluster data. We normalized the total number of binding events to the total region lengths and then calculated the enrichment Z-score for each factor.

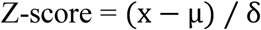

x: binding event counts of a factor in aCNEs, normalized by region lengths;

μ: mean of binding event counts of a factor in 1000 times permutation, normalized by region lengths;

δ: standard deviation of the binding event counts of a factor in 1000 times permutation.

For chromatin state enrichment analysis, the same strategy was used to determine chromatin states in randomly selected DHS regions. For each epigenome and each set of randomly selected regions, the percentage of regions showing “Active”, “Bivalent”, “Polycomb repressed”, “Quiescent” and “Other” states was calculated. The percentage distribution was used for calculate enrichment Z-score.

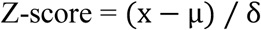

x: percentage of regions displaying a certain chromatin state in aCNEs
μ: mean of the percentages of regions displaying a certain chromatin state in 1000 times permutation
δ: standard deviation of the percentages of regions displaying a certain chromatin state in 1000 times permutation

### Processing and analyzing of published mouse cardiac TF ChIP-seq data

Mouse cardiac ChIP-seq or ChIP-exo data from the following studies were collected for annotating aCNEs.

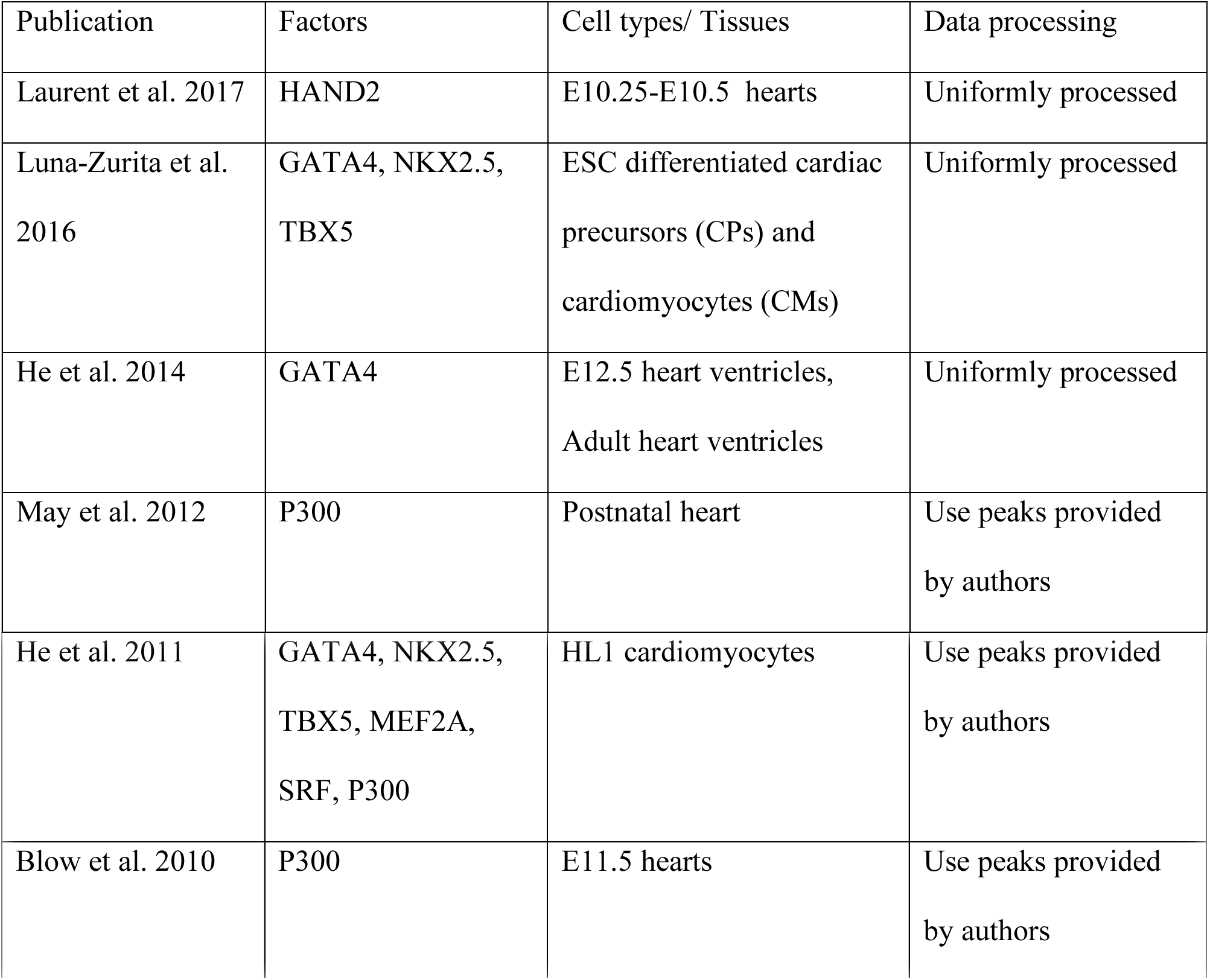

Data from studies of Laurent et al. 2017, Luna-Zurita et al. 2016 and He et al. 2014 were uniformly processed as below. Raw reads were trimmed with Trimmomatic (TRAILING:3 MINLEN:36) before align to mm9 genome assembly by BWA (version 0.7.8) under aln model ^91,98^. Reads with mapping quality score > 30 were kept for downstream analysis (samtools view - b -q 30). Replicates for the same sample were merged for peak calling by MACS2^99,100^ (version 2.7.9) (-q 0.01 -g 1.87e9). Peaks called by MACS2^99,100^ (Laurent et al. 2017, Luna-Zurita et al. 2016 and He et al. 2014) or provided by the authors (May et al. 2012, He et al. 2011 and Blow et al. 2010) were intersected with the mouse DHSs conserved with GFP+/- specific ATAC-seq peaks. If a cardiac TF or P300 peak overlaps a mouse DHS by at least one base pairs, this anciently conserved open region is considered as bound by this factor.

To generate tracks shown in Supplementary Fig. 1A, H3K27ac^19^ and cardiac transcription factor (GATA4, NKX2.5, TBX5)^22^ ChIP-seq data were used. Replicates for each factor each condition were merged by running Samtools^104^ (merge, Version 1.2) on the alignment files (in bam format) provided by the authors. Then BEDtools^103^ (genomecov, Version 2.19.1) were used to compute the reads coverage based on the merged bam files for track display.

### GATA motif scanning

6 Motifs preferred by GATA family TFs (MA0035.3 Gata1, MA0036.2 GATA2, MA0037.2 GATA3, MA0140.2 GATA1::TAL1, MA0482.1 Gata4 and MA0766.1 GATA5) were extracted from Jaspar 2016 core vertebrate motif database^105,106^ and scanned within the 162 GFP+ specific, human-zebrafish conserved ATAC-seq peaks by FIMO (version 4.12.0). If an ATAC-seq peak has a stretch of sequence that matches any of the 6 GATA motifs scanned (q < 0.3), this peak was counted as GATA motif positive. If different GATA motifs match to similar regions of a peak, it is only considered as one GATA motif occurrence in this peak due to the information redundancy within different GATA motifs. Only if multiple motifs hit non-overlapping regions of a peak will this peak be counted as having multiple GATA motifs. The 0.3 threshold of q score was determined by empirical evidence that the GATA motif mutated in experiments described in Supplementary Fig. 6 matches GATA5 motif with a q value of 0.291 in motif scanning. The full scanning results can be seen in Supplementary Table 5.

## Acknowledgements

We would like to thank all members of the Wilson and Scott Labs for helpful discussion and suggestions. Special thanks go to: Huayun Hou and Minggao Liang for guidance and comments on computational analyses; Angel Morley, Allen Ng, Scott Knox and Alejandro Salazar for expert fish husbandry; Michelle Ward and Sara Good for helpful comments of the manuscript; Michael Hiller for sharing the zCNEs full dataset and providing detailed explanation regarding the dataset. Caroline and Geoffrey Burns for sharing the *Tg(nkx2.5: ZsYellow) ^fb7^* transgenic line; Ryan Anderson for sharing the *Tg(βactin2:loxP-DsRed-STOP-loxP-EGFP)^s928Tg^* transgenic line; The SickKids-UHN Flow and Mass Cytometry Facility for FACS services; Sergio Pereira from The Center for Applied Genomics (TCAG) facility for conducting bulk and single-cell mRNA-seq; and ENCODE and Roadmap Epigenomics Mapping Consortium for generating and consolidating the open chromatin landscapes and chromatin state datasets. This research was supported by Hospital for Sick Children Restracomp Studentship (to XY) and Connaught International Scholarship (to XY); Ontario Trillium Scholarship (to MS); Heart and Stroke Foundation of Canada (to ICS and MDW, Grant-in-Aid G-16-00013798); NHBLI R01HL114948 (to BB); and the Natural Sciences and Engineering Research Council of Canada (NSERC) grant 436194-2013 (to MDW).

## Author Contributions

Conception and study design: XY, BB, ICS, MDW; Acquisition of data: XY, MS, PD; Analysis and interpretation of data: XY, ICS, MDW; supervision of work: MDW, ICS, and BB. XY, ICS, and MDW wrote the manuscript and all authors assisted with drafting and revision of the manuscript.

## Accession Codes

ATAC-seq and RNA-seq data are available at ArraryExpress under the accession number E-MTAB-6078 and E-MTAB-6077

